# Evolution induced state shifts in a long-term microbial community experiment

**DOI:** 10.1101/2025.10.10.680894

**Authors:** Mikko Kivikoski, Johannes Cairns, Shane Hogle, Sanna Pausio, Lutz Becks, Ville Mustonen, Teppo Hiltunen

**Affiliations:** Department of Computer Science, University of Helsinki, Finland; Organismal and Evolutionary Biology Research Programme, University of Helsinki, Finland; Department of Biology, University of Turku, Finland; Turku Collegium for Science, Medicine and Technology (TCSMT), University of Turku, Finland; Aquatic Ecology and Evolution, Limnological Institute, University of Konstanz, Germany

**Keywords:** *Aeromonas caviae*, antibiotic resistance, community dynamics, eco-evolutionary dynamics, experimental evolution, microbial community, state shift

## Abstract

Biological communities are complex, dynamic systems that underpin ecosystem functionality^1^, yet their long-term dynamics and predictability remain poorly understood^2^. Understanding how Darwinian evolution shapes these systems through eco-evolutionary feedbacks is a central challenge in ecology and evolution. Experimental studies using simplified microbial assemblages have yielded important insights into the ecological principles governing community states^3–5^. However, an important knowledge gap is how selection within member species drives changes of community state in multispecies systems. Here, we present a four-year evolution experiment involving a 23-species synthetic bacterial community propagated in two environments: a control medium and the same medium supplemented with the antibiotic streptomycin. Through combined analyses of community composition and genome evolution, we quantified the temporal changes in species abundances and the evolutionary trajectories of individual community members. The extended duration of the experiment enabled the detection of adaptive mutations and community state shifts that occur only over long evolutionary timescales. We show that community dynamics are environment dependent and reproducible across replicates, and that evolution of streptomycin resistance in a previously streptomycin-sensitive species on its own can induce abrupt community state shifts. Our results provide a direct demonstration of eco-evolutionary feedbacks within a multi-species community, revealing how a single adaptive mutation can reorganize complex ecological networks.

---

The biodiversity around us is an assembly of species that live and interact in communities. The composition of these communities – defined by the identities and abundances of their member species – emerges from assembly processes shaped by environmental conditions and interspecific interactions^6^. Classic examples show how environmental change and interactions shift communities, from state changes in lakes^7^ to microbiome restructuring after antibiotics^8^. Yet it remains unclear whether assembly follows general rules or reflects emergent, context-dependent properties^9,10^.

Evolution adds another layer of complexity, operating alongside ecological assembly to shape community trajectories. Microbes, with their short generation times, are particularly tractable models for studying these dynamics, although short-term studies often miss slower evolutionary changes. The long-term *E. coli* evolution experiment^11^ (LTEE) has shown that molecular evolution can persist for more than 60,000 generations, even after fitness increase plateaus. These insights have reshaped our understanding of adaptation and spurred theoretical advances such as clonal interference^12^ and predictive evolutionary frameworks^13– 15^. Yet single-species systems capture only part of the picture: real communities are far more complex. Ecological processes like species sorting structure community assembly and responses to perturbations^16–20^, but how evolution unfolds within these dynamics and feeds back to alter community states remains unresolved. Ecological conditions not only shape community structure but also drive Darwinian evolution, which can cascade back to reorganize entire communities^21^. These eco‐evolutionary feedbacks represent a key gap in our ability to predict long-term community outcomes^22^.

Bridging this gap requires closer integration of population genetics and community ecology^23,24^. Unlike single-species systems, multispecies communities involve direct interactions such as cross-feeding, toxins, and quorum sensing, as well as indirect interactions through shared-resource competition^25^. Theory and experiments increasingly show that the fitness effects of mutations depend on community context, producing evolutionary outcomes that diverge from those in isolated populations^26–30^. Understanding and predicting community dynamics will therefore require approaches that explicitly integrate ecological interactions with evolutionary change across timescales.

Extending long-term evolutionary approaches to multispecies microbial communities can illuminate how molecular evolution within one species triggers cascading effects that shift whole community states. Developing experimental frameworks that connect ecological and evolutionary processes over short and long timescales is a critical step toward predictive community ecology (Fig. 1). Here, we present our findings from the first 210 weeks of an ongoing community experiment involving a 23-species bacterial community. We tracked community composition, or the relative abundance of species, and the molecular evolution of member species in two environments with different levels of the antibiotic streptomycin. Six replicate communities were propagated in each environment. This design enabled us to examine the interaction between species sorting (ecological filtering) and evolutionary change in shaping community dynamics and to test whether molecular evolution influences the stability and trajectories of multispecies communities.

**Fig 1.**
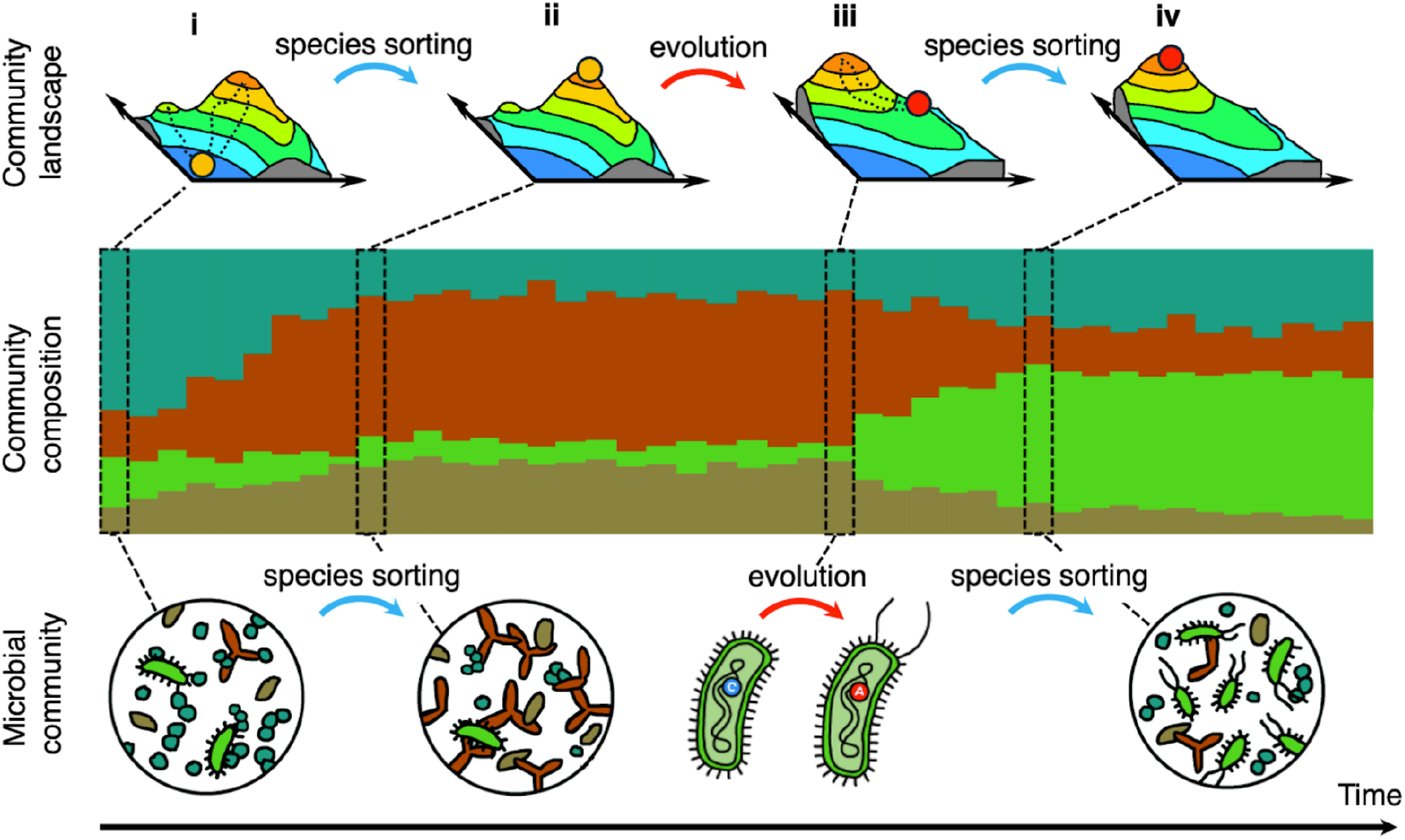
Microbial community composition, i.e. state, is shaped by ecological interactions, evolution, and their interplay. We studied how microbial communities (bottom) move across the community state space and change over time. The stacked bar chart (middle) shows the relative abundances of the member species (schematic). The top row depicts a schematic, low-dimensional, representation of the underlying high-dimensional dynamical system as a landscape where the microbial community (filled circle) moves due to ecological forces. The community moves typically towards one of the (local) maxima, and the succession path is dictated by the interaction of species with themselves and the environment and affected by stochastic forces such as demographic noise (panel **i**, dashed lines show example paths leading to different maxima). If the time-scale is long enough Darwinian evolution could change the dynamical multispecies system. Panel **iii**) demonstrates such a scenario where, after initial rapid species sorting (panel **ii**), a community member species acquires an adaptive mutation which alters the eco-environmental system and its dynamics, seeding a new high-dimensional dynamical system. Such a major event can be followed by a fast species sorting dynamics occurring again as the community moves towards a (possible) stable state (panel **iv**). Several, a priori potential dynamical scenarios could occur, and the figure illustrates one such scenario. For example, replicated microbial communities within an environment could go to different states, via different sequences of states, or to the same state via the same or different sequence of states, and these dynamics may or may not be affected by evolution.

Study environments, base (B) and streptomycin (S), had a substantial short-term effect on the community composition: the community states diverged markedly between absence and presence of streptomycin (Fig. 2). In both environments, a 16-days long serial transfer experiment, replicated 48 times, revealed rapid and highly repeatable species sorting, consistent with earlier studies (*e*.*g*. ref.^31^). These differences arose from eco-environmental effects, namely, differential sensitivity to streptomycin^32^ and the interactions among the species. The most pronounced difference in the community state was the relative abundance of *Aeromonas caviae* and *Citrobacter koseri*: *A. caviae* is sensitive to streptomycin and had the highest relative abundance in the base environment, but negligible abundance in streptomycin, whereas *C. koseri* which is resistant to streptomycin, showed the opposite pattern.

**Fig 2.**
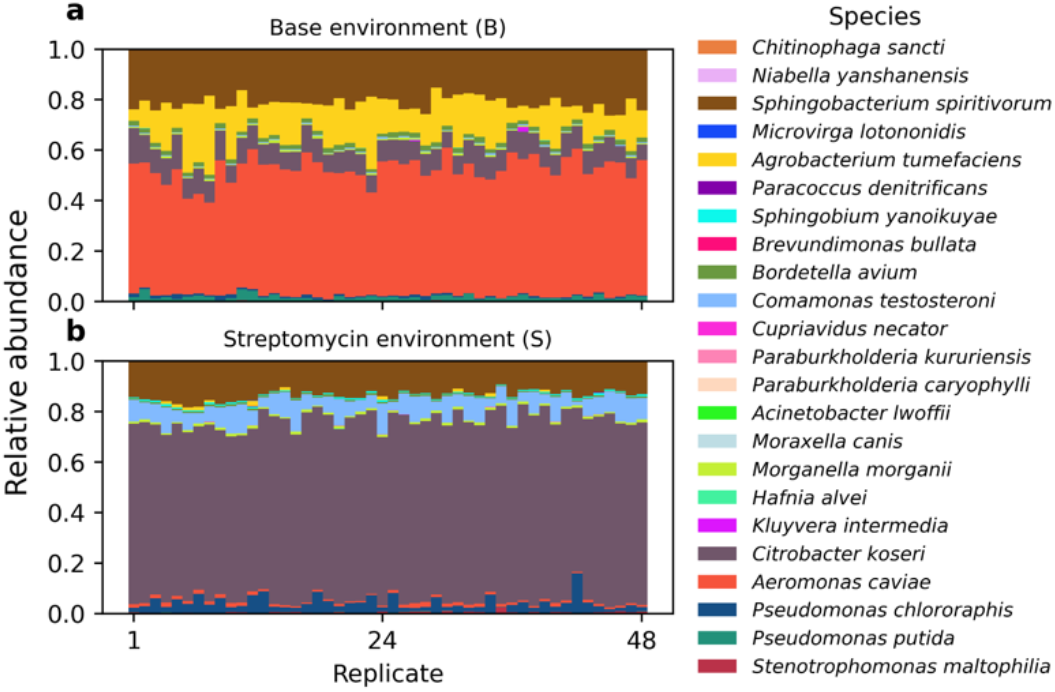
Community composition differs between (**a**) the base and (**b**) the streptomycin environments. Stacked bar charts show the relative abundances of the 23 species in 48 replicates after 16 days of serial transfer experiment.

The long-term experiment revealed successive, repeatable, community state transitions and diverging fates across the replicates. These dynamics were observed in a 210-week experiment with 48 longitudinal samples to assess the dynamics of community composition and evolution. The data revealed distinct succession in the two environments (Fig. 3). We segmented the time series into discrete epochs representing succession states. In both environments, except in replicate S6, the initial epoch was brief, representing a rapid transition from the inoculum to an eco-environmental determined state. In the base environment, all six replicates (B1–B6) followed a highly repeatable three-epoch succession pattern (Fig. 3; end-point repeatability index of 0.97 Fig. S1). In the streptomycin environment, five replicates (S1– S5) converged on a shared succession path (end-point repeatability index = 0.96, Fig. S1), although the number and timing of the epochs varied (Fig. 3). By contrast, replicate S6 diverged from the other replicates ultimately reaching a state resembling the base environment state (Fig. 3, end-point repeatability index of B1–B6 and S6 = 0.93, Fig. S1).

**Fig 3.**
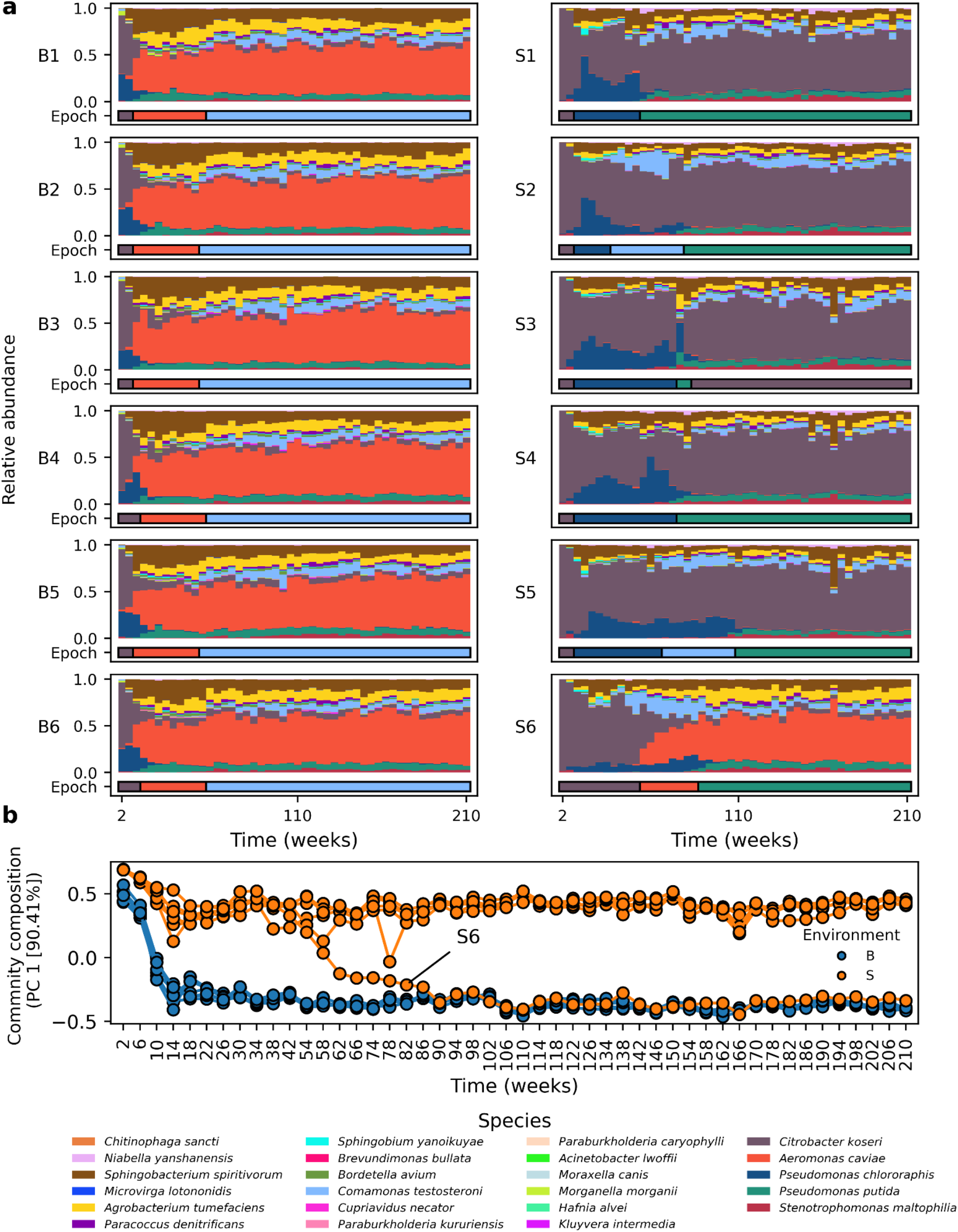
Community composition over time in six base and streptomycin environment replicates (B1–B6 and S1–S6, respectively). (**a**) Stacked bar charts show the relative abundances of each of the 23 species per time point. Horizontal bars show the epochs (3–4 per replicate) and their break points indicating state shifts. Epoch coloring indicates the species whose abundance change has the greatest impact on the inference of epoch change and differs from the previous epoch-determining species (see *Methods*). (**b**) Principal component 1 (PC 1) depicts the path of community succession. Each point corresponds to one timepoint in one replicate. Deviation of replicate S6 from the other S communities is highlighted. Percentage of the variance explained by PC 1 is indicated in the label of the y-axis.

An eco-evolutionary feedback drove the state shifts in the streptomycin environment. This feedback arose through selective sweeps of a known streptomycin resistance point mutation in the *rpsL* gene (Lys88Arg, *e*.*g*. ref.^33^)in *P. putida* (S1–S6) and *A. caviae* (S6) detected by metagenome sequencing and validated through clone isolation and growth assays. These adaptive events not only led to an increase in the species’ relative abundances (Fig. 4) but also reshaped the community structure: *P. chlororaphis* decreased as *P. putida* increased, and *A. caviae* increased at the cost of *C. koseri* (Fig. 3). Such shifts align with competitive exclusion among closely related taxa with similar metabolic pathways^32^. Together, the results show that evolutionary adaptation in *P. putida* and *A. caviae* reverberated through the community by restructuring the ecological interactions.

**Fig 4.**
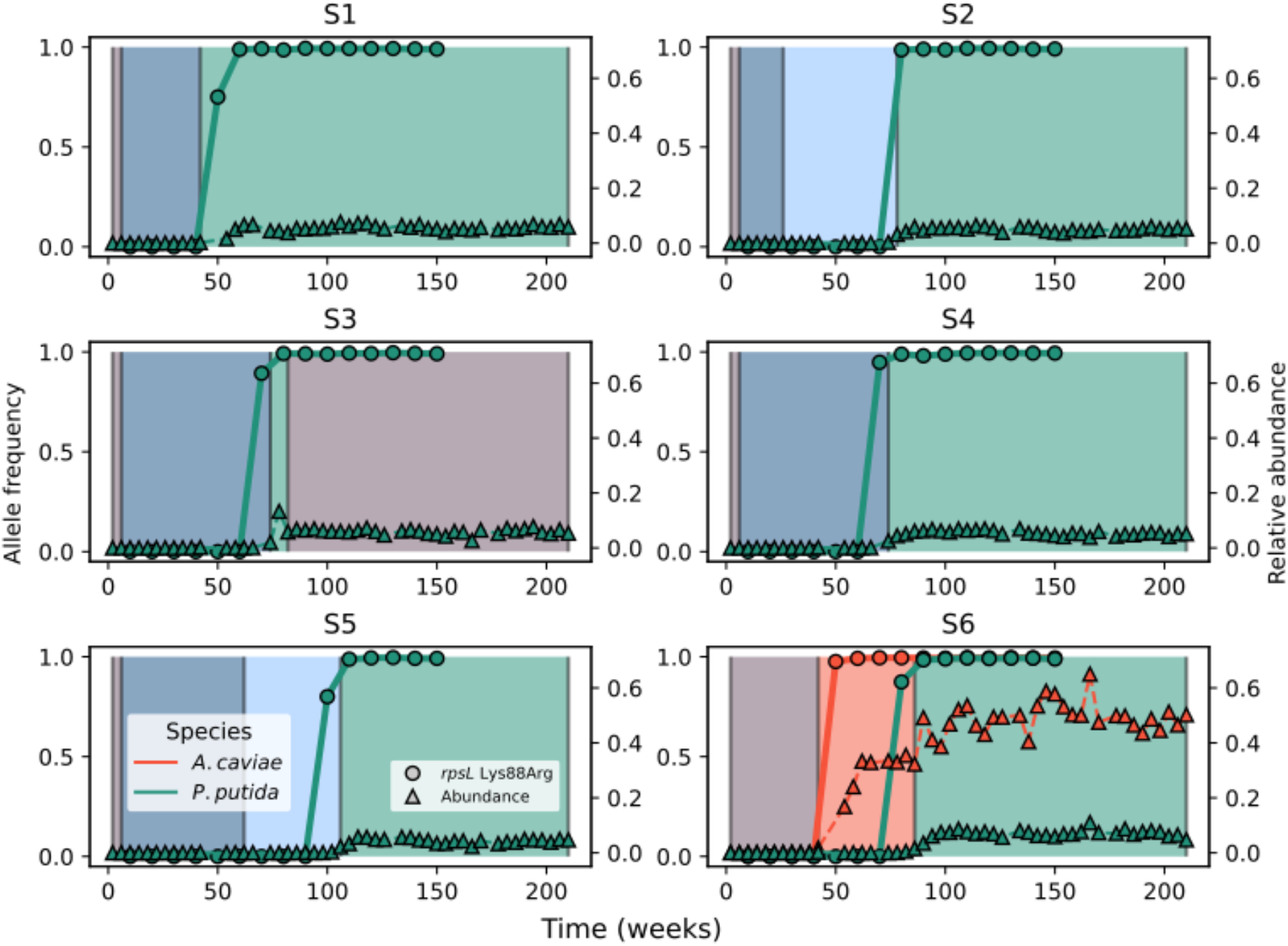
Selective sweeps of a non-synonymous point mutation (lysine to arginine in the 88th codon, Lys88Arg) in the gene *rpsL* in *Aeromonas caviae* (orange) and *Pseudomonas putida* (green) are linked to the community state shifts (*i*.*e*. epoch changes) in the streptomycin community replicates. Solid lines and circles show the allele frequency of the mutation (left y-axis) and the dashed lines with triangles show the relative abundance of the species (right y-axis). The background shows the epochs of Fig. 3.

Introducing resistant clones of *P. putida* and *A. caviae* to the ancestral communities recapitulated the state shifts observed in the long-term experiment. This outcome aligned with time-resolved community and genomic data (Figs. 3, 4) which indicated that streptomycin resistance evolution in *P. putida* drove the state shift observed in all six replicates, while that of *A. caviae* explained the unique trajectory of community S6. To test causality, we introduced sensitive and resistant *P. putida* or *A. caviae* clones (isolated from the week 151) to the ancestral communities growing either in the base or streptomycin environment. Resistant clones of both species established successfully and altered community composition whereas the sensitive clones did not establish in the streptomycin environment (Figs. S2, S3). Especially in the streptomycin environment, resistant *A. caviae* shifted the community composition toward that of the base environment (repeatability index = 0.92, Figs. S2, S3), confirming its role as the driver of the community composition. We next asked whether the long-term communities would respond similarly. After 207 weeks, resistant *A. caviae* was introduced to the streptomycin communities and monitored for seven weeks. Invasion succeeded in S1–S5 (in S6 it was already present) shifting community compositions toward the S6 state (Fig. 5).

**Fig 5.**
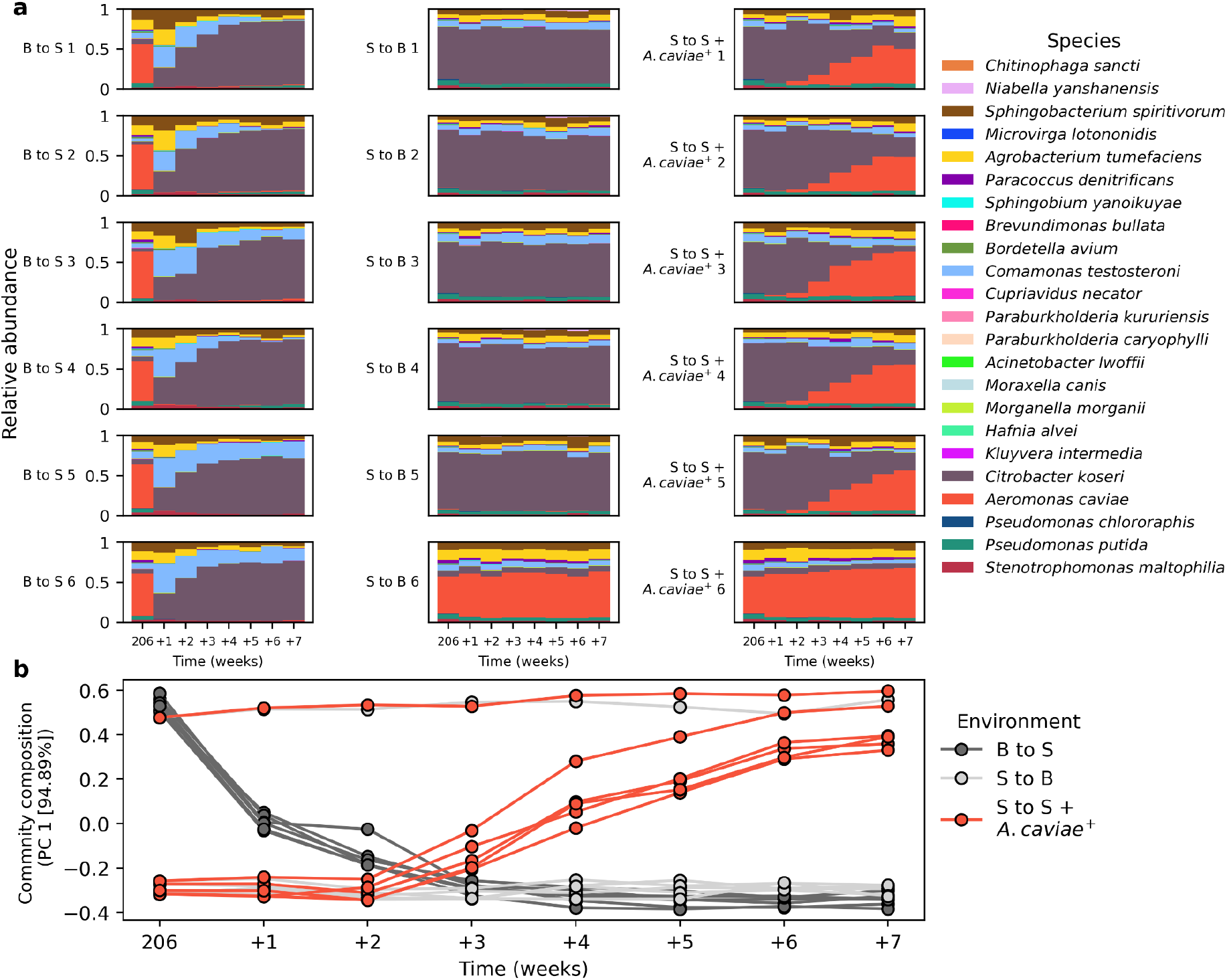
Consequences of changing the environment (from base, B to streptomycin, S or *vice versa*) or introducing a resistant *A. caviae* clone in a streptomycin environment after an evolutionary history of 207 weeks (see *Methods*). (**a**) The bar charts show the community composition from week 206 in the main experiment to week seven in the experiment. (**b**) The scatter plot shows the community composition (principal component 1) as a function of time. Each point corresponds to one timepoint in one replicate. Changing the environment of the B communities to streptomycin changed the community structure while the opposite, changing S communities to no streptomycin, did not change the community structure. Resistant *A. caviae* (*A. caviae*^+^) was able to invade communities S1–S5 where it was absent.

Switching the environments demonstrates that state shifts induced by resistance evolution were robust. We tested this by reversing the environments after week 207. Adding streptomycin to base environment communities B1–B6 eradicated *P. putida* and *A. caviae* while favoring *C. koseri* (Fig. 5). This response arose from the streptomycin sensitivity of *P. putida* and *A. caviae*, and the resistance of *C. koseri*. The similarity of outcomes across the replicates supports the conclusion that long-term base communities had converged to highly repeatable states.

By contrast, removing streptomycin from streptomycin adapted communities S1–S5 left their composition unchanged (Fig. 5). In particular, *A. caviae* did not reappear in communities S1– S5 where it was absent or had a very low abundance (below detection threshold) after the streptomycin treatment. Finally, removing streptomycin had no detectable effect on the relative abundance of *A. caviae* in the S6 community or *P. putida* in S1–S6 communities, suggesting that any fitness cost of streptomycin resistance was absent or mitigated. Together these findings show that eco-evolutionary dynamics can entrench microbial communities in stable states that persist even under reversal of the environment.

In this study, the environment (base versus streptomycin) determined the long-term community state (Fig. 3). Trajectories showed a rapid early transition followed by extended periods of stability, reminiscent of single-species LTEE^34,35^. Unexpectedly, in the streptomycin treatment, it was not the antibiotic stress alone but the subsequent evolution of resistance in *A. caviae* and *P. putida* over one to two years that drove the transition to new community states (Fig. 4). Once resistance arose, communities followed either one distinct trajectory (S1– S5) or one resembling the base environment (S6). These states persisted even after antibiotic removal, indicating resilience, and could be reproducibly induced by introducing a resistant *A. caviae* clone (Fig. 5). The response of community composition to the appearance of resistant *A. caviae* was independent of the community history. This suggests that the uniqueness of *A. caviae* resistance evolution in S6 was not due to that particular community but rather because of its very low population size which limits the chances for mutations. *A. caviae* is more sensitive to streptomycin than *P. putida*, which explains the higher rate of resistance mutations in *P. putida*^*32*^. Although additional resistance mutations were detected in other species (Table S1), they did not appear to shape community-level dynamics. Evolution was also prevalent in the base environment, with selective sweeps in *Stenotrophomonas maltophilia, Comamonas testosteroni*, and *Paracoccus denitrificans* (Fig. S4). However, the timing and length of epochs across base replicates were highly consistent, suggesting that while evolution was common, it caused the community trajectory to diverge from repeatable patterns only with resistance mutations in key taxa. This shows that streptomycin exerts a strong species-specific control on the community and once species resolve the stress they revert the community composition to the non-perturbed state reminiscent of evolutionary rescue.

Linking community ecology with population genetics has long been a theoretical goal, yet empirical connections remain scarce. The studies examining the effect of community context on evolution have revealed broad patterns, with mechanisms and species driving changes remaining elusive, *e*.*g*. ref.^21,36,37^. Here, we directly tested how ecological and evolutionary change interact to shape community dynamics and stability. We show that a *de novo* mutation in a single key species can redirect the ecological state of an entire microbial community – a striking demonstration of eco-evolutionary feedback dynamics in communities. Contrary to the expectation that antimicrobial resistance carries substantial fitness costs^38^, resistant *A. caviae* and *P. putida* produced community-level outcomes comparable to those without streptomycin. This pattern may reflect strong interspecific differences in competitive ability compared to those within species (sensitive versus resistant subclones) or low fitness costs of *rpsL* mutations^39^. More broadly, our results demonstrate that evolutionary innovation can reshape community trajectories and reinforce resilience, highlighting how eco-evolutionary dynamics may be harnessed to better understand, predict, and eventually manage the stability of microbial communities in natural and applied settings.

## Materials and Methods

### Bacteria

The 23 bacterial species forming the synthetic community were from the HAMBI Culture Collection (University of Helsinki). The species were chosen for the community, based on the previous research demonstrating their ability to grow together in mesocosm studies^17,18,32^. The bacterial species used were: *Acinetobacter lwoffii, Aeromonas caviae, Agrobacterium tumefaciens, Brevundimonas bullata, Bordetella avium, Chitinophaga sancti, Citrobacter koseri, Comamonas testosteroni, Cupriavidus necator, Hafnia alvei, Kluyvera intermedia, Microvirga lotononidis, Moraxella canis, Morganella morganii, Niabella yanshanensis, Paraburkholderia caryophylli, Paraburkholderia kururiensis, Paracoccus denitrificans, Pseudomonas chlororaphis, Pseudomonas putida, Sphingobacterium spiritivorum, Sphingobium yanoikuyae, Stenotrophomonas maltophilia*.

### Long-term serial passage experiment

Bacterial strains were cultured in 25 ml microcosm flasks that contained 6 ml of 20% R2A medium (Neogen). The experiment consisted of two conditions: base environment (no streptomycin) and streptomycin. The streptomycin concentration in the culture medium was 20 μg × ml^-1^ of streptomycin sulfate and we report the concentration as the concentration of the sulfate. There were six replicate communities in both environments. The experiment was initiated by adding 20 μl mixture of the 23 bacteria species (in equal proportions) to the medium. The flasks were incubated in a shaking cabin at the temperature of 25 C for seven days. After seven days, 500 μl of the medium was transferred to a fresh medium as described above. The culture remaining after the seven day growth cycle was processed as follows: 100 μl culture was collected for measuring optical density (OD) with Absorbance 96 plate reader (Byonoy), two 900 μl culture specimens were pelleted and stored at -20 C for DNA extraction and sequencing, and 500 μl of culture was frozen with 85 % glycerol (1:1) for further use (*e*.*g*. clone isolation). As part of the transfer protocol, the community was reseeded with a tiny amount (less than 1% of the total community biomass) of the ancestral community every sixth transfer. This was done in order to avoid stochastic extinctions during the experiment and to allow evolution *via* mutations and migration of those species that went extinct or were present at very low abundance.

### Short-term serial passage experiment

The long-term serial passage experiment demonstrated evolution of streptomycin resistance in *Pseudomonas putida* and *Aeromonas caviae*, and community composition changes related to this. These dynamics took years to observe and to assess their repeatability, we performed a 16-days long experiment with a transfer at every 24 hours. The same ancestral community, culture media and streptomycin concentrations as in the long-term experiment were used. The communities were cultured on 96-well plates in 1 ml volume. As a control, the ancestral 23-species community was grown in streptomycin and streptomycin-free conditions (48 replicates of both conditions). To assess community dynamics related to streptomycin resistance, resistant or sensitive *A. caviae* and/or *P. putida* clones, isolated from the long-term experiment (see the section below), were introduced to the community. Introduction was done at the beginning of the experiment or such that one species was introduced at the beginning and the other one at the middle of experiment. The introduction assay was performed in a full-factorial setting involving both streptomycin concentrations, resistant and sensitive clones of both species added at the beginning or middle of the experiment.

### *Aeromonas caviae* and *Pseudomonas putida* clone isolation

Streptomycin sensitive and resistant *A. caviae* and *P. putida* clones were isolated from the 151st transfer of the long-term experiment. The putatively sensitive clones were isolated from the base replicates and the putatively resistant clones were isolated from the streptomycin replicates. Resistant *A. caviae* clones were present only in replicate S6. The sensitivity to streptomycin was studied by culturing each clone in an array of streptomycin concentrations (0, 1.25, 2.5, 5, 10, 20, 40, 80, 160 μg × ml^-1^). Finally, each selected clone was whole-genome sequenced to confirm that they carry the same streptomycin resistance mutation as observed in the long-term experiment. For the sequencing of the clones, DNA was extracted with DNeasy 96 Blood & Tissue Kit (Qiagen). Sequencing was carried out at SeqCenter (Pittsburgh, Pennsylvania, USA) with Illumina NovaSeq X Plus sequencer. The sensitive clones were selected from those that did grow in absence of streptomycin but not in the concentration of 20 μg × ml^-1^, and the resistant clones were selected from those that did grow in absence of streptomycin and in the concentration of 20 μg × ml^-1^.

### Validation with the evolved communities

The short-term serial passage experiment could validate the findings in the long-term experiment, namely, that streptomycin affects the community composition and that resistance of *A. caviae* or *P. putida* changes it. However, it remains open how a change in the streptomycin condition would affect the long-evolved communities, and, if a resistant *A. caviae* could invade those five streptomycin communities where it did not evolve the streptomycin resistant. To address these questions we made an additional transfer at week 207: each replicate was transferred to a fresh culture medium as usual but an additional transfer was made to the opposite environment, *i*.*e*. from base to streptomycin and *vice versa*. This additional community was transferred weekly according to the normal transfer protocol and followed for eight weeks by analysing the community composition every week. The second line of validation was to introduce a resistant *A. caviae* clone to the S1–S6 communities to address the question if *A. caviae* would become the dominant species also in those communities where it did not evolve resistance during the study period. This was carried out by introducing one of the resistant *A. caviae* clones used in the other validation experiment to each streptomycin community. These communities were also followed for eight weeks. As a control of the experiment communities were also maintained in their environment and a sensitive *A. caviae* clone was introduced to S1–S6 communities (Fig. S5). Community composition throughout this experiment was assessed with 16S rRNA amplicon sequencing similar to the main study (see below). The 8th timepoint of the experiment had data with suspected errors and it was excluded from analyses. In addition, the 3rd time point in S to S replicate 6 was excluded due to the low number of sequencing reads (Fig. S5).

### 16S rRNA amplicon sequencing and community composition analyses

Relative abundances of the 23 species were based on 16S rRNA amplicon sequencing. Genomic DNA for community composition analysis was extracted from the frozen sample of every 4th transfer using the DNeasy 96 Blood & Tissue Kit (Qiagen). Sample quality and quantity was assessed using NanoDrop (Thermo Scientific) and Quant-iT 1X dsDNA HS kit (Invitrogen). Amplicon library was constructed with an in-house library preparation method. Briefly, V3 region of 16S gene was amplified and combinatorially indexed using tailed primers, purified with NGS Normalization 96-Well Kit (Norgen Biotek) and subsequently amplified and dual indexed with primers compatible with Illumina platforms^40^. Finally, the reaction was purified with Sera-Mag particles (Cytiva) and the quality of the library was evaluated using Bioanalyzer High sensitivity DNA kit (Agilent). Sequencing was carried out at the Finnish Functional Genomics Centre, Turku, Finland.

Analysis of the relative abundances of the 23 species was based on mapping the paired-end 16S rRNA amplicon reads to references of the 23 species 16S rRNA genes following the protocol used by Cairns *et al*.^*18*^. In brief the analysis protocol begins with quality trimming, merging and filtering of the paired-end 16S reads. Then the reads are mapped to a reference genome of the 16S rRNA genes from the 23 species^41^. Finally, the species counts are normalized by species-specific 16S rRNA gene copy number to obtain relative abundances. See gitlab.utu.fi/slhogl/hambiAmplicon for the documentation of the workflow.

At the beginning the longitudinal 16S data started from the second week and lasted until the 210th transfer, yielding 53 timepoints (one per every fourth week). Of the 53 timepoints, three had missing data (weeks 46, 50, and 70) in multiple samples, and week 174 had clearly deviating data in multiple samples. Week 130 sample was sequenced in a different laboratory and slightly different protocol, yielding qualitatively but not quantitatively comparable data. These five time points were excluded and as a result the longitudinal community composition data consisted of 48 timepoints.

### Segmentation of the community composition into epochs

The longitudinal data on the community composition was segmented in order to understand state shifts and succession of the communities. Segmentation was done with an R^42^ package *dpseg* (ver. 0.1.1, ref.^43^). The function dpseg in the package was modified to allow segments to have a minimum length of two timepoints (lower limit originally three). Here, *Rcpp* package (ver. 1.0.14, ref.^44,45^) was utilised. The scoring matrix for the segmentation was calculated by taking the average community composition for every 2, 3, 4…48 timepoint long segments and summing up the Kullback-Leibler divergence of the segment average to each time point within the segment. We hereby call the segments as epochs.

### Repeatability of the community composition

Repeatability of community composition among replicates was quantified with repeatability index as described by Venkataram and Kryazhimskiy^2^. Community similarity metric *s* between communities *i* and *j* at time *t* was calculated as 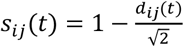, where *d*_*ij*_(*t*) is Euclidean distance of the 23 elements long community composition vectors at time *t*. Repeatability indices for the compared replicates were calculated as the mean of the similarity metrics per every time point.

### Metagenome sequencing and analyses

Evolution, *i*.*e*. emergence of *de novo* mutations and changes in their frequency was studied with whole-genome sequencing of the metagenome, *i*.*e*. the total DNA of the community. For this, DNA was extracted from the sample of every 10th transfer, spanning the weeks 10–150, yielding 15 timepoints in total. DNA extraction was carried out with DNeasy 96 Blood & Tissue Kit (Qiagen), in a 96-well plate format. The concentration and fragmentation of the extracted DNA was inspected with Quant-iT 1X dsDNA HS kit (Invitrogen) and NanoDrop ND-1000 (Thermo Scientific). Extractions of 60–400 ng of DNA at 25–40 μl volume were taken to the sequencing facility (Finnish Functional Genomics Centre, Turku, Finland) for specimen preparation and sequencing.

Metagenome sequencing was carried out as paired-end whole-genome sequencing (2 x 150 bp reads). Whole-genome sequencing was done with NovaSeq 6000 S4 technology (version 1.5, Illumina). Adapter trimming was executed at the sequencing facility with bcl2fastq2 conversion software (Illumina). The total number of sequencing reads per sample was ca. 20 million reads.

Prior to read mapping and other bioinformatic analyses the technical quality of the metagenomic sequencing data was assessed with FASTQC software (v. 0.11.9 www.bioinformatics.babraham.ac.uk/projects/fastqc/). None of the performed tests indicated any technical problems that would have led to discarding of the data.

Read mapping was done with the BWA-mem algorithm^46^. All samples were mapped against a meta reference, constructed from the high-quality reference genomes of each 23 species^41^ and a ciliate *Tetrahymena thermophila* strain SB210^47^. *T. thermophila* was included in the meta reference in order to map possible *Tetrahymena* reads present in other samples of the sequencing batch. Mapped reads were further processed with SAMtools (v. 1.21, ref.^48^) methods sort, fixmate, markdup and merge, respectively, to prepare alignments for variant calling.

Variant calling was executed per replicate community, providing alignments of each timepoint as one sample. The workflow and analysis steps followed those in Hoffmann *et al*.^*26*^. Short variants, *i*.*e*. single-nucleotide variants (SNV), multi-nucleotide variants (MNV) and short indels, were called with GATK (v. 4.5.0.0, ref.^49^) tools MUTECT2 and FilterMutectCalls. In FilterMutectCalls, microbial mode (--microbial-mode) was applied. SNVs with filter quality ‘PASS’ were used in further analyses. After this step, clusters of SNVs were identified with GATK tool VariantFiltration, by defining the number of SNVs that make up a cluster to two (--cluster-size 2) and size of the search window to 35 bp (--cluster-window-size 35). Ten base pair long flanks were added to each SNV to make a list of SNV cluster regions with Bedtools. VariantFiltration was further used to exclude these regions and putative mobile elements from the list of SNVs. MNVs and indels with filter quality ‘PASS’ and those that were only observations within a 35 bp long window were retained. Finally the effects of SNVs, MNVs and indels with a minimum mapping depth of 5, were predicted with snpEff (v. 4.3t, ref.^50^) based on the gene annotations from Cairns *et al*.^*18*^. Mutations in protein coding genes, annotated as ‘CDS’, were used in the analysis of evolutionary changes.

## Analysis and interpretation

Downstream analyses and visualization of the data, *e*.*g*. community composition and identified mutations were carried out in Python 3.10 environment using the libraries Numpy (v. 1.26.4, ref.^51^), Scipy (v. 1.13.0, ref.^52^), Seaborn (0.13.2, ref.^53^), pandas (v. 2.2.2, ref.^54,55^), Matplotlib (v. 3.8.4, ref.^56^), and Scikit-learn (v. 1.4.2, ref.^57^).

## Acknowledgements

We want to acknowledge Meri Lindqvist and Katja Salminen at the Center of Evolutionary Applications (University of Turku), Elisa Zakharova, Ida-Marija Hyvönen, Beda Anttila, Tiia Korpi, Olli Pitkänen, Inga-Katariina Aapalampi, Milla Similä, Atte Aapalampi, Ida Aarni, Trine Link, Emmi Hirvonen, Julia Saloranta, Hilla Pellikka, Linda Nevala, Fanny Koskela, and Irma Warinowski for contribution to the laboratory work. Anthony Sun, Eva Kisdi and Seppe Kuehn are thanked for discussions. Mikhail Shubin is thanked for his assistance with drawing figure 1. This work was funded in part by the Research Council of Finland (Multidisciplinary Center of Excellence in Antimicrobial Resistance Research, grant #346126; grants #330886 & #327741 to TH; grants #346128 and #364234 to VM). This study was supported by Finnish Functional Genomics Centre, University of Turku and Åbo Akademi and Biocenter Finland. The authors wish to acknowledge CSC – IT Center for Science, Finland, for computational resources.

## Author contributions

Conceptualization, MK, LB, VM, TH

Formal analysis, MK

Investigation, all authors

Methodology, MK, JC, SP, SH

Experimental work, SP

Writing – original draft, MK, JC, LB, VM, TH.

Writing – review & editing, all authors

Funding acquisition, VM, TH

Supervision, VM, TH

VM and TH equal contribution.

