## Supplementary figures and tables for "Evolution induced state shifts in a long-term microbial community experiment"

#### Supplementary material for the manuscript
#### Evolution induced state shifts in a long-term microbial community experiment

Mikko Kivikoski[1,2], Johannes Cairns[1,2,3,4], Shane Hogle[3], Sanna Pausio[3], Lutz Becks[5], Ville Mustonen[1,2,*,†], Teppo Hiltunen[3,*,†]

Affiliations
1: Department of Computer Science, University of Helsinki, Finland
2: Organismal and Evolutionary Biology Research Programme, University of Helsinki, Finland
3: Department of Biology, University of Turku, Finland
4: Turku Collegium for Science, Medicine and Technology (TCSMT), University of Turku, Finland
5: Aquatic Ecology and Evolution, Limnological Institute, University of Konstanz, Germany

[*]Corresponding authors: Teppo Hiltunen, Ville Mustonen
[†] Equal contribution

### Table of contents

##### Supplementary Figures

Fig. S1
Fig. S2
Fig. S3
Fig. S4
Fig. S5

##### Supplementary Tables

Table S1

### Supplementary Figures

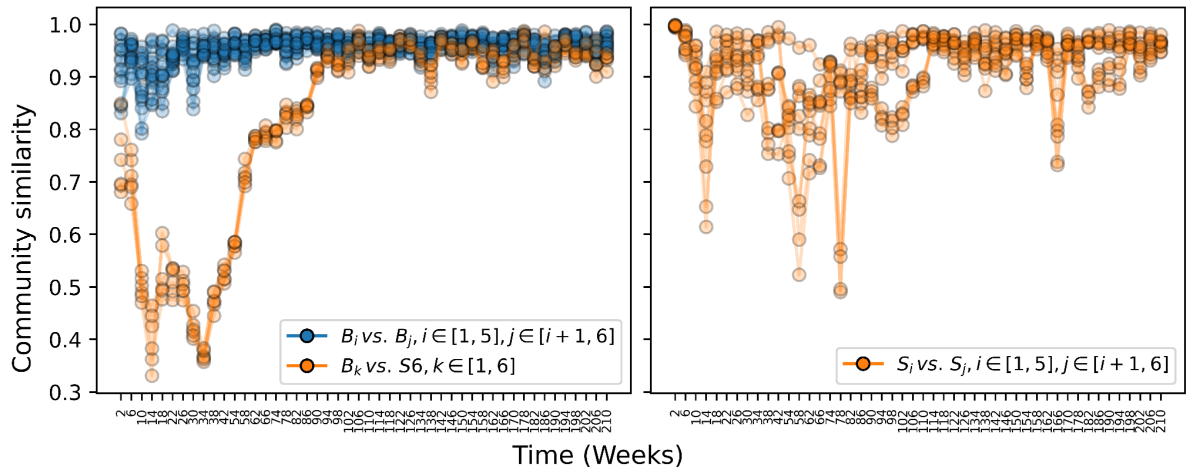


**Fig. S1**

Community similarity across time in all pairwise comparisons of B1–B6 and S6 communities (left panel) and S1–S5 communities (right panel). The community state repeatability indices at the end of the experiment were 0.97, 0.93, 0.96 among the B1–B6 communities (blue lines in the left panel), B1–B6 communities and S6 community (orange lines in the left panel) and among S1–S5 communities (orange lines in the right panel), respectively.

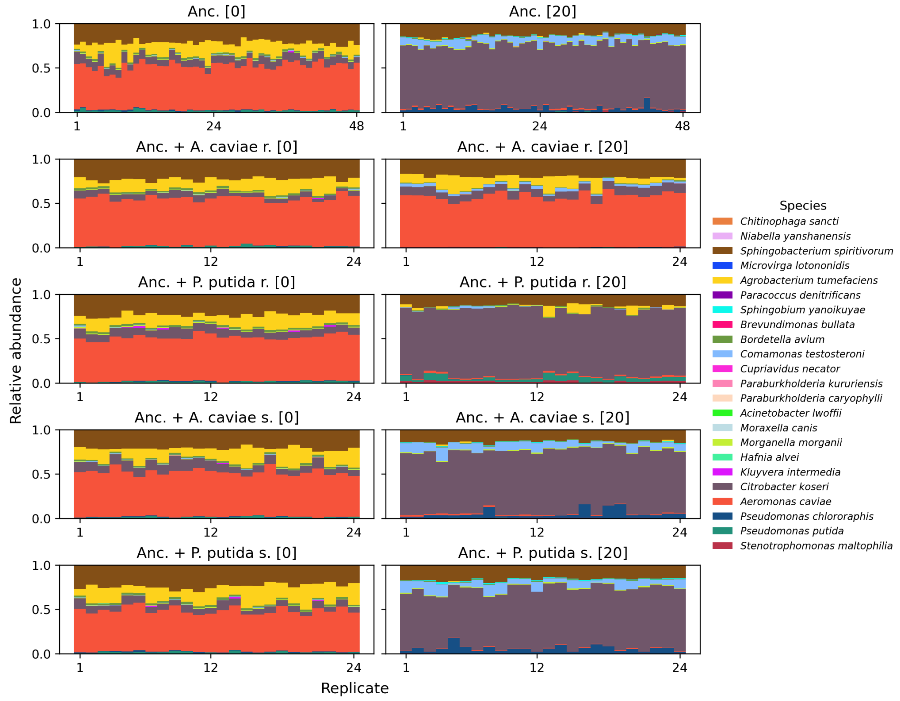


**Fig. S2**

Community composition after a 16-day serial transfer experiment in the base and streptomycin environment (left and right panel, respectively). Stacked bar charts show the relative abundances of the 23 species in 24 or 48 replicates. Top panels show the ancestral community (Anc.) and the panels below show those with streptomycin sensitive (s.) or resistant (r.) *A. caviae* or *P. putida* clone.

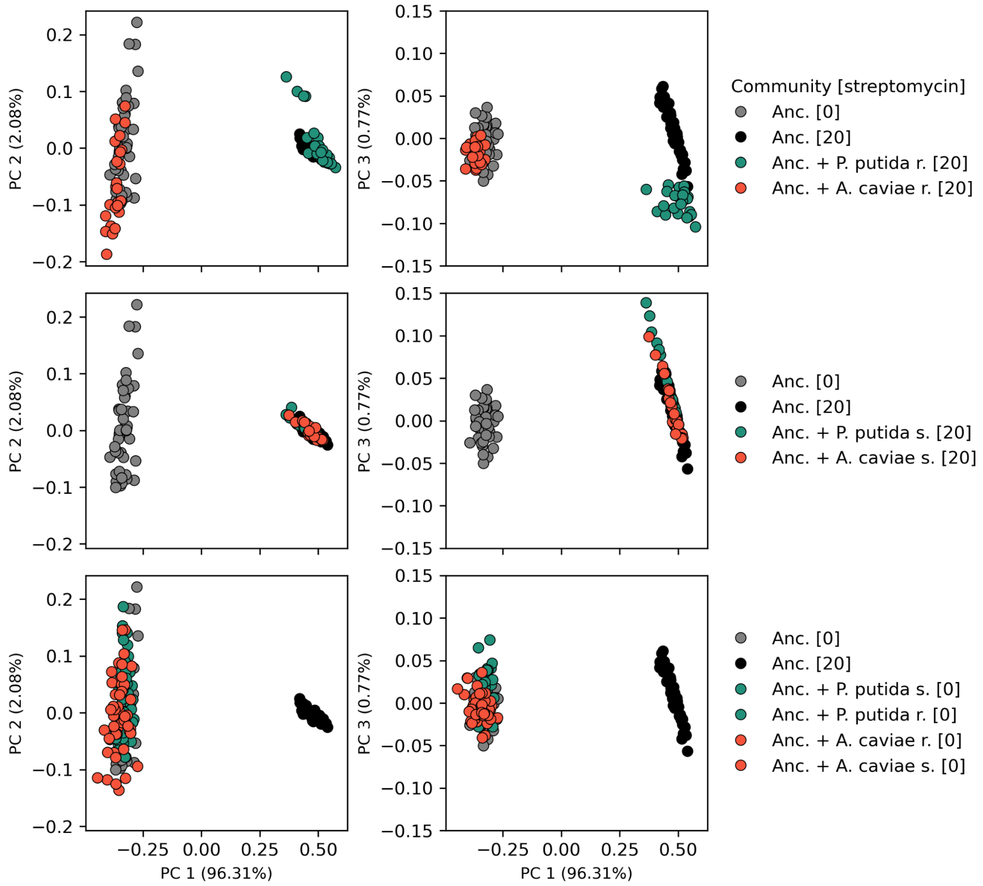


**Fig. S3**

Principal component analysis of the community composition after a 16-day serial transfer experiment in the base and streptomycin environment (left and right panel, respectively). Grey and black dots show the ancestral community (Anc.) replicates in base and streptomycin environments, respectively. The orange and green dots show the ancestral community with an introduced *A. caviae* or *P. putida* clone, respectively.

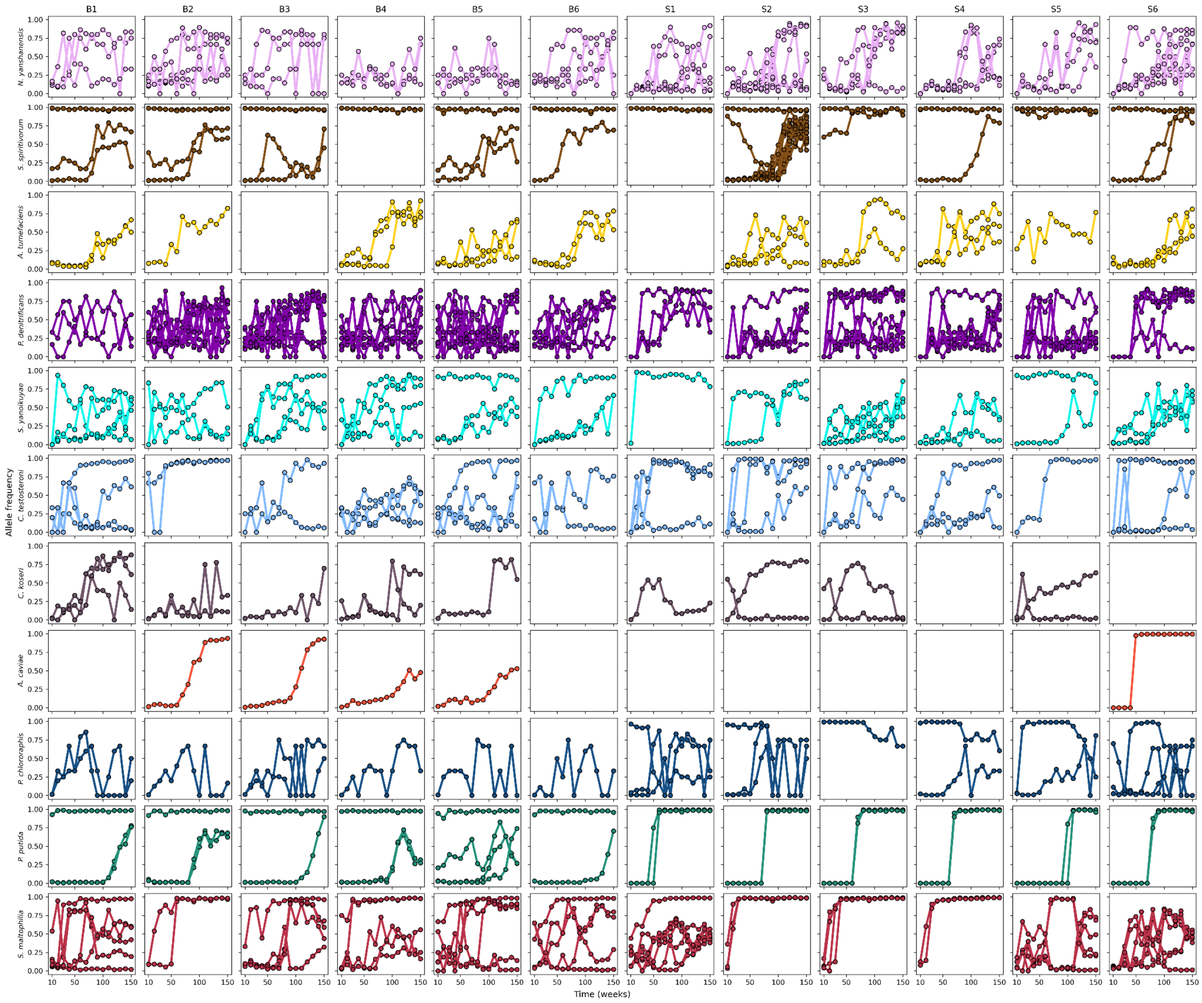


**Fig. S4**

Allele frequencies across time for mutations reaching allele frequency of 0.5. The 11 species with minimum relative abundance of 0.05 or more are shown.

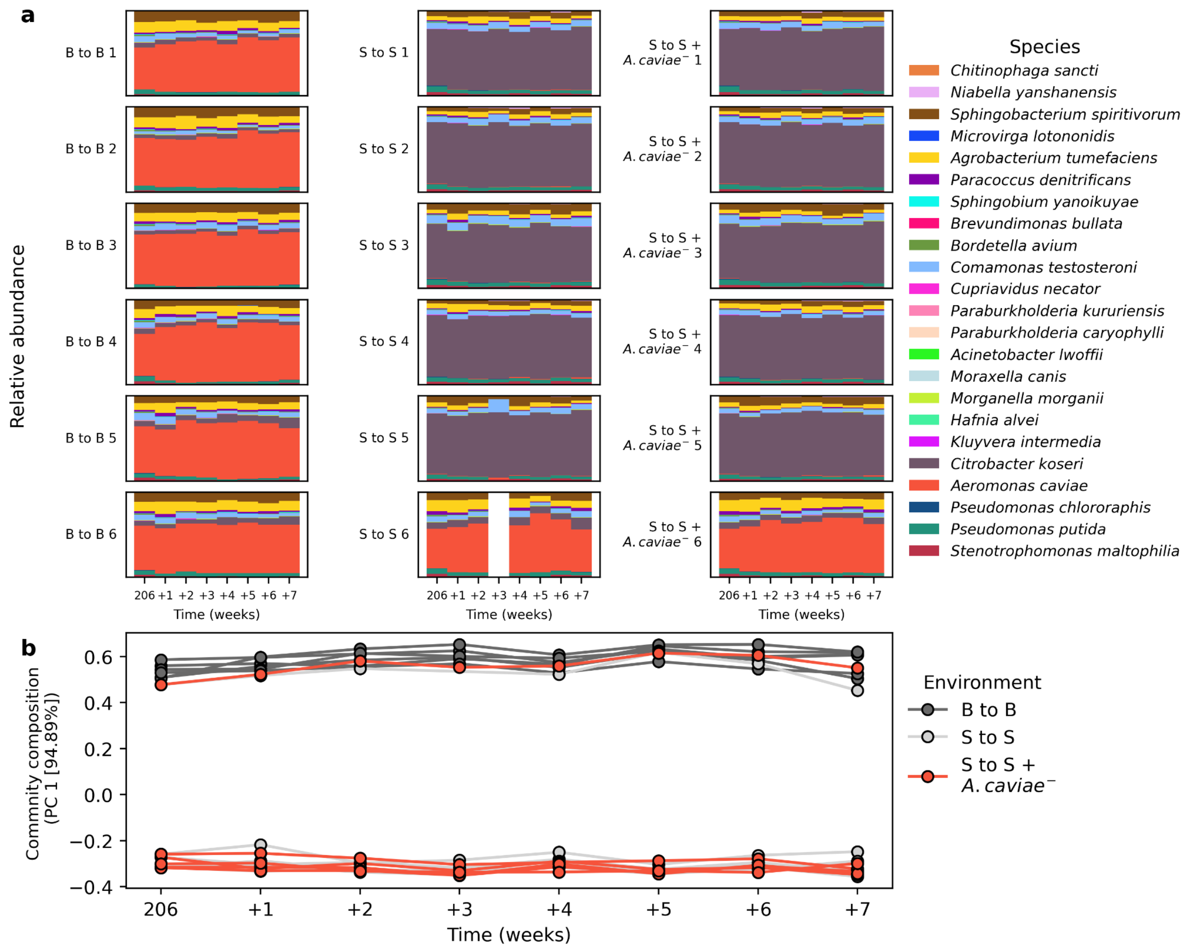


**Fig. S5**

Control treatments for the validation experiment. Communities from B and S environments maintained in the same environment (B to B and S to S, respectively), and communities in the S environment with an introduced streptomycin sensitive *A. caviae* clone (S to S + *A. caviae*$^{-}$). The missing data point in the S to S replicate 6 at week 3 has technically erroneous data.

### Supplementary Tables

**Table S1**

**Recurrently mutated genes (intragenic mutation in more than one replicate) with mutations reaching high allele frequency (> 0.80) at any point of the long-term experiment. Species with relative abundance above 0.05 at any point of the study are shown, except *C. koseri* which did not have any mutations matching these criteria. Genes *rsmG and rpsL* with streptomycin resistance causing mutations are highlighted in bold.**

| Species | Genes | Replicates mutated |
|---|---|---|
| *Aeromonas caviae* | *fliI*, ***rpsL*** | B2; S6 |
| *Agrobacterium tumefaciens* | *ropA* | B2, S3, S4, S6 |
| *Comamonas testosteroni* | *flhA, Outer membrane porin protein 32,* ***rsmG*** | B3, S1; S1, S2; S6 |
| *Niabella yanshanensis* | *clpX, rapA, TonB* | B1, B3; S4, S5; B1, B2, S6 |
| *Paracoccus denitrificans* | *dctP, gatB, pseI,* ***rpsL***, ***rsmG***, *rutR, tam* | B1, B4; S3, S6; B3, S3, S5, S6; S5; S1–S6; B2–B6, S1, S6; B5, B6 |
| *Pseudomonas chlororaphis* | ***rsmG*** | S1–S6 |
| *Pseudomonas putida* | ***rpsL*** | S1–S6 |
| *Sphingobacterium spiritivorum* | ***rpsL***, ***rsmG***, *TonB* | B1–B6, S1–S6; S2, S3, S5; S2 |
| *Sphingobium yanoikuyae* | *btuB, PKHD-type hydroxylase* | S3, S6; B1, B3–B6, S2 |
| *Stenotrophomonas maltophilia* | *btuB, cnrA, pglF, smf* | B1, B2, B4–B6, S2, S3, S5; B1, B3, B5; B1, B2, B6, S2, S6; B1, B3–B6, S1 – S3 |
